# Causal, Predictive or Observational? Different Understandings of Key Event Relationships for Adverse Outcome Pathways and Their Implications on Practice

**DOI:** 10.1101/2024.06.25.599864

**Authors:** Zheng Zhou, Jeroen L.A. Pennings, Ullrika Sahlin

**Affiliations:** Center for Environmental and Climate Science, Lund University, Sweden; National Institute for Public Health and the Environment, the Netherlands

## Abstract

The Adverse Outcome Pathways (AOPs) framework is pivotal in toxicology, but the terminology describing Key Event Relationships (KERs) varies within AOP guidelines. This study examined the usage of causal, observational and predictive terms in AOP documentation and their adaptation in AOP development. A literature search and text analysis of key AOP guidance documents revealed nuanced usage of these terms, with KERs often described as both causal and predictive. The adaptation of terminology varies across AOP development stages. Evaluation of KER causality often relies targeted blocking experiments and weight-of-evidence assessments in the putative and qualitative stages. Our findings highlight a potential mismatch between terminology in guidelines and methodologies in practice, particularly in inferring causality from predictive models. We argue for careful consideration of terms like causal and essential to facilitate interdisciplinary communication. Furthermore, integrating known causality into quantitative AOP models remains a challenge.

## 1 INTRODUCTION

### 1.1 Background

Regulatory toxicology and environmental risk assessment in the twenty-first century need to handle the assessment of ever-increasing numbers of chemicals, and at the same time adhere to the 3R principles to reduce, refine and replace animal experiments (Pallocca et al. 2022). The Adverse Outcome Pathways (AOPs) methodology facilitates the extrapolation of evidence at lower biological levels from animal and non-animal methods for the inference of adverse outcomes at higher biological levels (Villeneuve et al. 2014a). An AOP is defined as an evolving conceptual framework to organize existing knowledge pertaining to a molecular initiating event (MIE), an apical adverse outcome of interest (termed adverse outcome; AO), key measurable/ observable events (termed key events; KEs), and the linkages among MIE, AO and KEs (termed key event relationships; KERs) (Villeneuve et al. 2014a). A list of definitions and abbreviations is presented in the supplementary information (SI). The robustness and reliability of the extrapolation across events at different biological levels therefore depends on the characterization and inference of KERs (Villeneuve et al. 2024), which could be causal, observational and predictive.

### 1.2 Causal Inference and Its Challenges in Key Event Relationship Evaluation

Causal inference pursues the unbiased understanding of how one condition, the cause, affects another condition, the outcome, free from the threats to internal validity (Guzelian et al. 2005). Randomized experiments are the gold standard for causal inference (Guzelian et al. 2005; James et al. 2015) as they can approximate unbiased counterfactual estimates of treatment effects (Höfler 2005; Pearl 2009). Toxicological studies typically implement randomized experiments in controlled laboratory conditions, which reveal unbiased estimates of the association between the randomized assignment (e.g. dose) to the compound of interest and the experimental outcome (e.g. response), thereby establishing a causal relationship(Höfler 2005). The mechanism of randomized controlled trials is reviewed in the SI.

When randomized experiments are not feasible, inference could rely on observational data. Quasi-experimental methods involve selecting subgroups from observational data to approximate unbiased average treatment effects, assuming specific conditions are met (Angrist and Pischke 2009). These methods have been employed in various fields such as economy, social science, psychology and epidemiology (Angrist and Pischke 2009; Bagiella et al. 2015; Fréchette et al. 2022). Observational data can also be evaluated using evidence-based approaches. The Bradford Hill considerations (Hill 1965), an early evidence-based framework for causal inference on observational, epidemiological data, has been modified to infer causal estimates in various contexts (Guzelian et al. 2005). For example, the evolved, rank-ordered Hill considerations for mode of action human relevance framework can evaluate the weights of evidence supporting hypothesized KERs in modes of action (MOAs) (Meek et al. 2014). However, neither quasi-experimental methods nor evidence-based frameworks fully guarantee unbiased estimates of causality. Therefore, results based on observational data have lower reliability compared to those from randomized experiments, which are termed as “causal estimates” in fields such as econometrics and social science. Within this article, we use the term “observational” to distinguish causal estimates from counterfactual, causal findings, emphasizing the observational nature of the data.

Despite the involvement of experimental toxicological data, there could be challenges to obtain the empirical evidence needed for causal inference of KERs (Spînu et al. 2022). The association between the dose and an outcome is causal, while the association between two outcomes (KEs) is not necessarily causal, due to the inability to randomize upstream KEs. For example, consider two KEs observed from the same dose-response experiment, with a hypothetical KER between them (Figure 1). This experiment generates two pairs of dose-response data, one for *KE*_*up*_ and one for *KE*_*down*_ respectively. While this setup provides unbiased estimates of the dose effects on *KE*_*up*_ and *KE*_*down*_ (Figure 1 left panel), *KE*_*up*_ itself is the consequence of the treatment and thus not randomized among subjects. The lack of randomization in *KE*_*up*_ implies that the estimates of the KER may not be causal (Figure 1 right panel). Direct evaluation of observed changes in *KE*_*down*_ conditional on *KE*_*up*_ is vulnerable to the threats to internal validity (Lipton and Ødegaard 2005), such as omitted variable bias from confounding. An instance of confounding bias is that, even if *KE*_*up*_ does not actually cause change in *KE*_*down*_, simultaneous changes in both KEs might still occur due to a third, unobserved confounding event. The relationship between *KE*_*up*_, *KE*_*down*_ and a potential confounder differs from the concept of non-adjacent KERs described in the handbook (Villeneuve et al. 2024). This is because a confounder might affect both *KE*_*up*_ and *KE*_*down*_ while a non-adjacent KER is when a third KE, which may not affect *KE*_*up*_, is bridged over to *KE*_*down*_. The estimated effect of *KE*_*up*_ on *KE*_*down*_ would be biased without isolating the confounding bias. For example, overdose of acetaminophen can cause hepatotoxicity (Hinson et al. 2010) and nephrotoxicity (Mazer and Perrone 2008). Both liver and kidney damage are suggested to be mediated by the depletion of glutathione (Jaeschke and McGill 2015; Mazer and Perrone 2008). The normal function of glutathione is involved in the detoxification of N-acetyl-p-benzoquinone imine, a toxic metabolite of acetaminophen (Casarett 2008). If the role of glutathione depletion is not properly accounted for, it could introduce confounding bias to falsely suggest hepatotoxicity leads to nephrotoxicity upon acetaminophen overdose.

**Figure 1:**
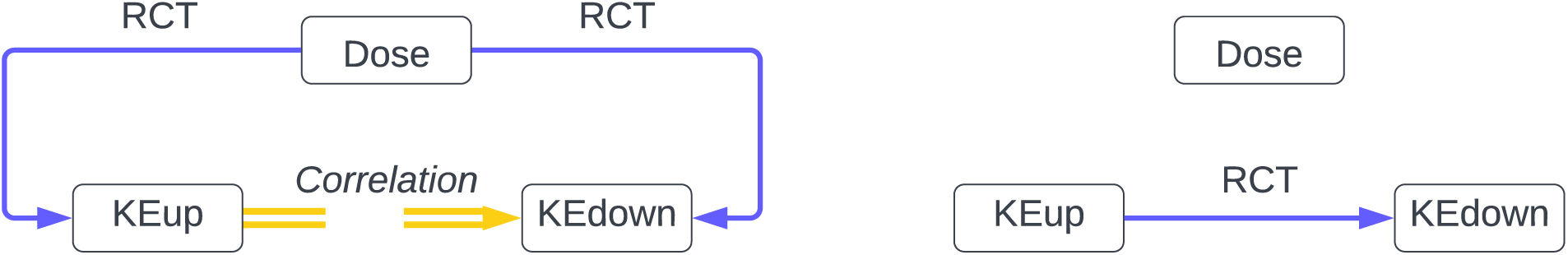
Directed acyclic graph of key event relationship evaluation. RCT: randomized controlled trials. Left shows that treatment effects are causal on both key events respectively; right shows what is needed for a causal evaluation

### 1.3 Distinguishing Causal and Predictive Inference under the Context of Adverse Outcome Pathways

Predictive models have been used in the application of traditional mechanistic study of toxicology (Helma 2005), such as physiologically based pharmacokinetic models (Sager et al. 2015). It also contributes to the development and application of alternatives to animal methods (Khabib et al. 2022), such as quantitative structure activity relationships (Barber et al. 2024; OECD 2014) and quantitative system toxicology (Bloomingdale et al. 2017).

The difference between causal and predictive inference has led to debate and confusion (Young 2019). Causal inference falls under the framework of treatment effect evaluation (Young 2019), whose goal is to approximate unbiasedness (Young 2019) using randomized experiments and, when those are not possible, non-experimental methods (quasi-experimental (Angrist and Pischke 2009), evidence-based frameworks). In contrast, the objective of predictive inference is to make accurate predictions. The term “predictive” reflects the scope of utility and application, and falls outside the treatment effect framework (Young 2019). Predictions can be accurate even when biased by confounders (Pearl 2009). For example, social-economic status is recognized as a major confounder to the adverse neurological effects of lead exposure during early childhood (Bellinger et al. 1992). However, some early studies accurately predicted neurological deficits based on dentine lead levels among children without adjusting for social-economic status (Herbert et al. 1979).

The difference between “causal” and “predictive” terms can be further understood in relation to mechanism and quantification. Mechanism implies that a causal relationship defines a directed linkage from the cause to the outcome (Russo and Williamson 2007), whereas the opposite is the error of reverse-causality. There is also a temporal requirement in a causal relationship that the cause must occur prior to the outcome, as emphasized through various iterations of Hill considerations (Becker et al. 2015; Cox 2018; Hill 1965; Meek et al. 2014). In contrast, a predictive relationship can be non-directional, i.e., knowing the temporal order between events is not necessary for prediction.

Quantification implies that a predictive relationship is validated on a strong statistical strength of association. In contrast, a causal relationship does not necessarily include a description of the magnitude of causality. Two events with a causal relationship could be even statistically independent of each other, i.e., zero correlation. We present two examples in the supplementary information for these two points respectively.

Furthermore, practical differences between predictive and the treatment effect frameworks exist in many commonly used methods. Consider a linear regression with multiple covariates, whose parameters are optimized using maximum likelihood estimation. This regression can be used for both prediction and causal inference (Young 2019). When this regression is used as a predictive model, the goal is to improve the model’s predictive performance (Young 2019), through the selection and optimization of the design matrix. When this regression is used as a causal model, bias adjustment is performed on the data prior to model construction, such as control selection (Cinelli et al. 2022) and backdoor elimination (Pearl 2009). The goal is to use the adjusted data and identify predictors that have significant coefficients (p values using maximum likelihood estimation and null hypothesis testing) (Angrist and Pischke 2009). The same model works differently for prediction and treatment effect evaluation. On the one hand, bias adjustment on data may not improve predictive performance (Brookhart et al. 2010; Davies et al. 2013). Including predictors of local significance may not improve global performance (Obermeyer and Emanuel 2016). On the other hand, optimizing the design matrix can not reduce the bias without prior bias adjustment (Chen et al. 2021).

The distinction between causal, observational and predictive inference is particularly relevant to context of toxicology and AOP development. This difference should be considered in the assembly of KEs and KERs. Traditionally, KEs and KERs are characterized among measured/ observed events from existing data, which is retrospective and evidence-based. In contrast, recent *in silico* predictive models enable prospective forecast of KEs and KERs for new compounds with minimal structural analysis (Hemmerich and Ecker 2020). The distinction between causal, observational and predictive inference should also be considered taking the difference between MOAs and AOPs. A major distinction between MOAs and AOPs is that MOAs are chemical specific while AOPs are not (Villeneuve et al. 2014a; Villeneuve et al. 2024). Evaluation of KERs under the MOA framework (Boobis et al. 2006) involves the evolved MOA Hill considerations (Meek et al. 2014), where the confidence in KER causality is crucial along the MOA of the compound of interest. In the context of AOP development, it has been suggested that the causality is necessary between the AO and compounds on all AOP models including predictive ones (Spînu et al. 2022).

However, it remains unclear whether it is necessary to require causality in all KERs in a chemical agnostic AOP. Given the chemical agnostic principle of AOPs, the supporting data for AOP development may include a wide range of compounds, making it common to observe varying KE activities across compounds. Therefore, it is crucial to consider the appropriate application of causal, observational and predictive inference in AOP development.

### 1.4 Study Objectives

Given the difference between the terms “causal”, “observational” and “predictive” and its relevance to KER characterization, we hypothesize that there could be misuse of the terms and it hinders the effective evaluation of KERs. To test this hypothesis, the objectives of this study are to: 1) examine the use of terminology describing KERs across AOP documentations and 2) investigate the adaptation of different terms in the development and application of AOPs. Achieving these objectives may facilitate a continuous development of AOP guidelines and a pragmatic implementation of AOP framework in academic, regulatory and industrial contexts.

## 2 METHODS

### 2.1 Terms for Key Event Relationships

We performed a text analysis on two AOP documentations focused on setting the definitions, principles and guidelines for AOP development: AOP developer’s handbook (Villeneuve et al. 2024) (abbreviated as the handbook) and OECD Guidance Document for the Use of Adverse Outcome Pathways in Developing Integrated Approaches to Testing and Assessment (OECD 2017) (abbreviated as OECD guidance thereafter). For a comprehensive analysis on the descriptions of KERs, manual searches were performed in the handbook and OECD guidance with the following keywords: “key event relationship” OR “KER” OR ((relationship OR link) AND “key event”). We also noted the sentences before and after the located keywords for context.

### 2.2 Adaptation of terminology

We searched journal articles published by May 2024 to identify how different terminology has been adapted in development and application of AOPs. Our search was performed using online scholarly databases such as Google Scholar and PubMed. We gave preferences to sources from the “Animal-free Safety assessment of chemicals: Project cluster for Implementation of novel Strategies” (ASPIS) research cluster, including: ONTOX (ONTOX 2024), RISK-HUNT3R (RISK-HUNT3R 2024) and PrecisionTox (PrecisionTox 2024). We prioritized information from ASPIS due to the alignment between this article’s emphasis on AOPs and ASPIS’s perspective on non-animal methodologies. These searches aimed to identify exemplary cases to illustrate typical scenarios where the terms describing KERs have been adapted, rather than to conduct a systematic review of all relevant literature.

The identified information was organized by how the method of adaptation fits into the stages of AOP development and quantitative AOP (qAOP). AOP development consists of three operationally defined stages (OECD 2017; Villeneuve et al. 2014a). In contrast, there is no established guidance or consensus on the definition or stages of qAOP. Therefore, acting phases for qAOP application were based on pragmatic objectives from published literatures (Conolly et al. 2017; Perkins et al. 2019; Spinu et al. 2020), including prediction of AO (Jeong et al. 2018), derivation of point of departure (POD) (Perkins et al. 2019), and optimization of testing strategies (OECD 2017; Perkins et al. 2019).

## 3 RESULTS

### 3.1 Usage of terminology

KER descriptions in the OECD guidance are primarily based on the AOP developer’s handbook, with minor editorial changes for the use in integrated testing strategies. Therefore, results of our text analysis focus on KER descriptions identified from the handbook, highlighting the text examples located from the sections where the definitions and principles of AOP development are set. Results show that with a clear distinction between the terms “causal” and “predictive”, the handbook’s usage of them varies across sections (Table 1). For instance, in the handbook Section 3 KER Description on page 25, KERs are described in different ways: as predictive *“Within the AOP framework, the predictive relationships that facilitate extrapolation are represented by the KERs”* or as causal *“KERs are always described in the form of a directed relationship (one-way arrow) linking an upstream “causing” event to a downstream “responding” event”*. In some instances KERs are described with both “causal” and “predictive”, e.g., on page 6 Table 1 Definitions of Key Terms that *“A scientifically-based relationship that connects one key event to another, defines a causal and predictive relationship between the upstream and downstream event”*.

**Table 1:**
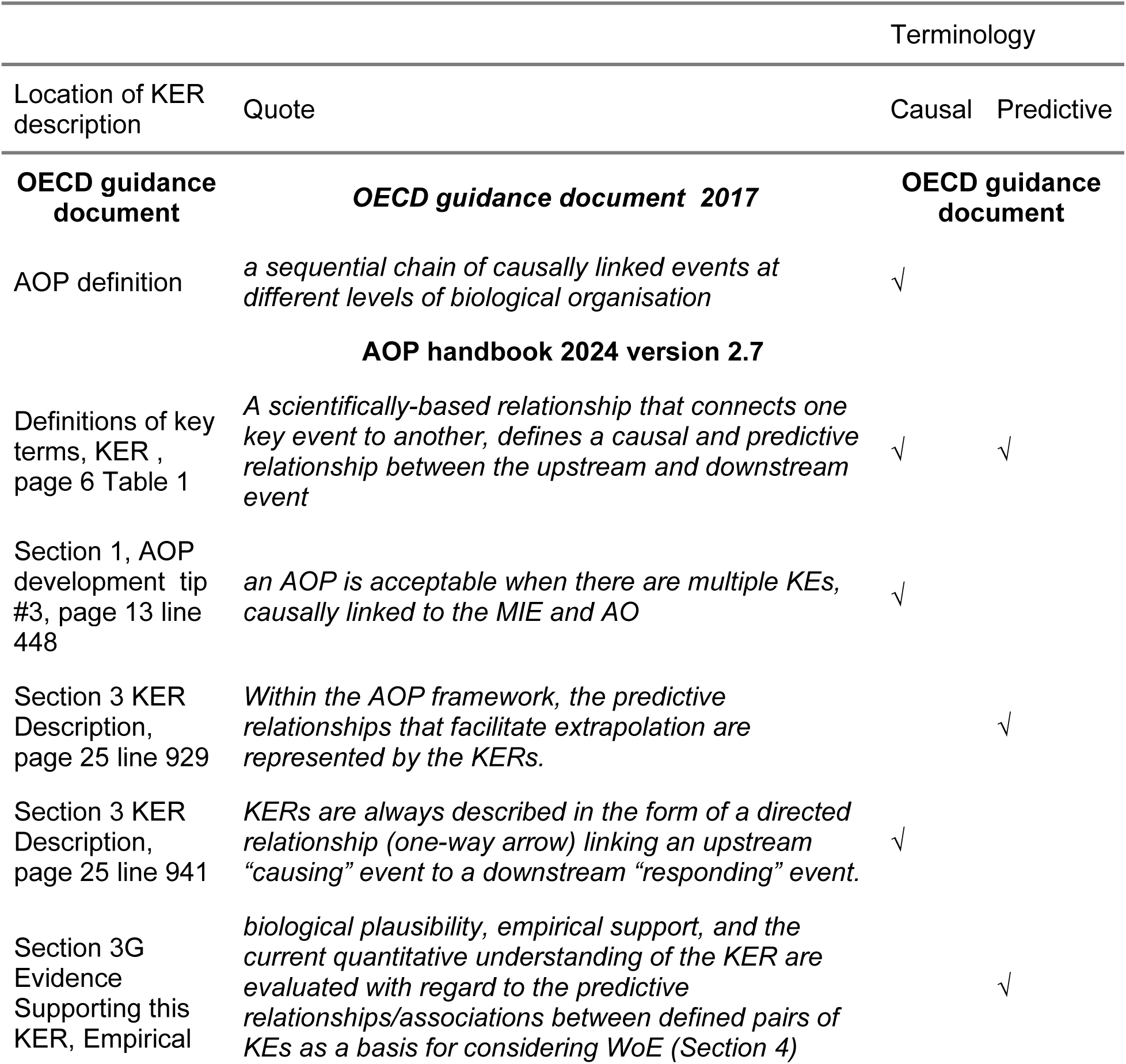
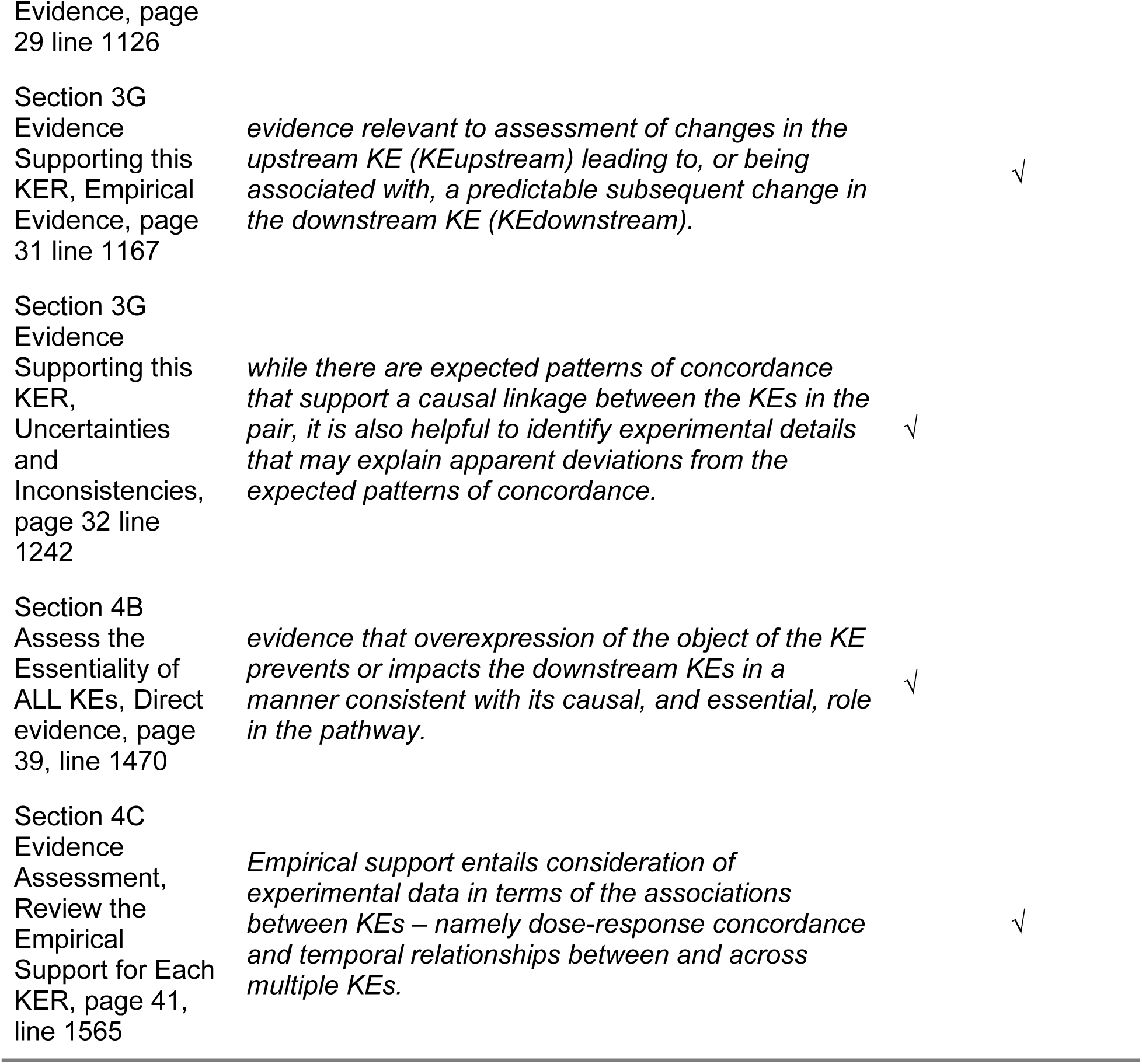
Occurrence of terminology describing KERs in two AOP documentations.

Compared to instances where “causal” or “predictive” are used separately, the inclusion of both terms suggests the consideration of dual requirements in KER description. The dual requirements do not necessarily imply a conflict against separate use of terms. When both causal and predictive inference are needed for KER description, one or both may be emphasized at different stages of AOP development.

The dual requirements are also evident in the handbook’s protocol for evaluating KER supporting evidence. The handbook evaluates KER supporting evidence (Section 3G) through 1) biological plausibility, 2) empirical evidence and 3) current quantitative understandings. The handbook’s guidance to assess biological plausibility (annex 1) is adapted from the evolved MOA Hill considerations (Meek et al. 2014), which is observational as mentioned in the Introduction of this paper. Empirical evidence is assessed (handbook page 31 subsection ii) for predictive utility that *“assessment of changes in the upstream KE (KEupstream) leading to, or being associated with, a predictable subsequent change in the downstream KE (KEdownstream)”*. Moreover, within the same section, the uncertainty and inconsistency of empirical evidence is evaluated that *“while there are expected patterns of concordance that support a causal linkage between the KEs in the pair, it is also helpful to identify experimental details that may explain apparent deviations from the expected patterns of concordance”*. The handbook’s evaluation of current quantitative understandings of KERs, considering the weights of empiricial evidence supporting KER causality, is dedicated to prediction *“to define the precision with which the state of the downstream KE can be predicted from knowledge of the state of the upstream KE”* (page 34). Therefore, it seems that the handbook sets the definitions, descriptions and protocols for evaluation of KERs based on a dual focus of causal and predictive inference. The term “observational” or the more commonly recognized “causal estimates” does not appear in the handbook, nor the OECD guidance.

### 3.2 Adaptation of terminology in AOP development and qAOP application

The handbook’s dual requirement on both causal and predictive KERs provides a comprehensive framework for AOP development. Examples identified from published literature are categorized by which term among “causal”, “observational” and “predictive” was adapted, and displayed throughout stages (Villeneuve et al. 2014a) of putative, qualitative and quantitative AOP development (Table 2).

**Table 2:**
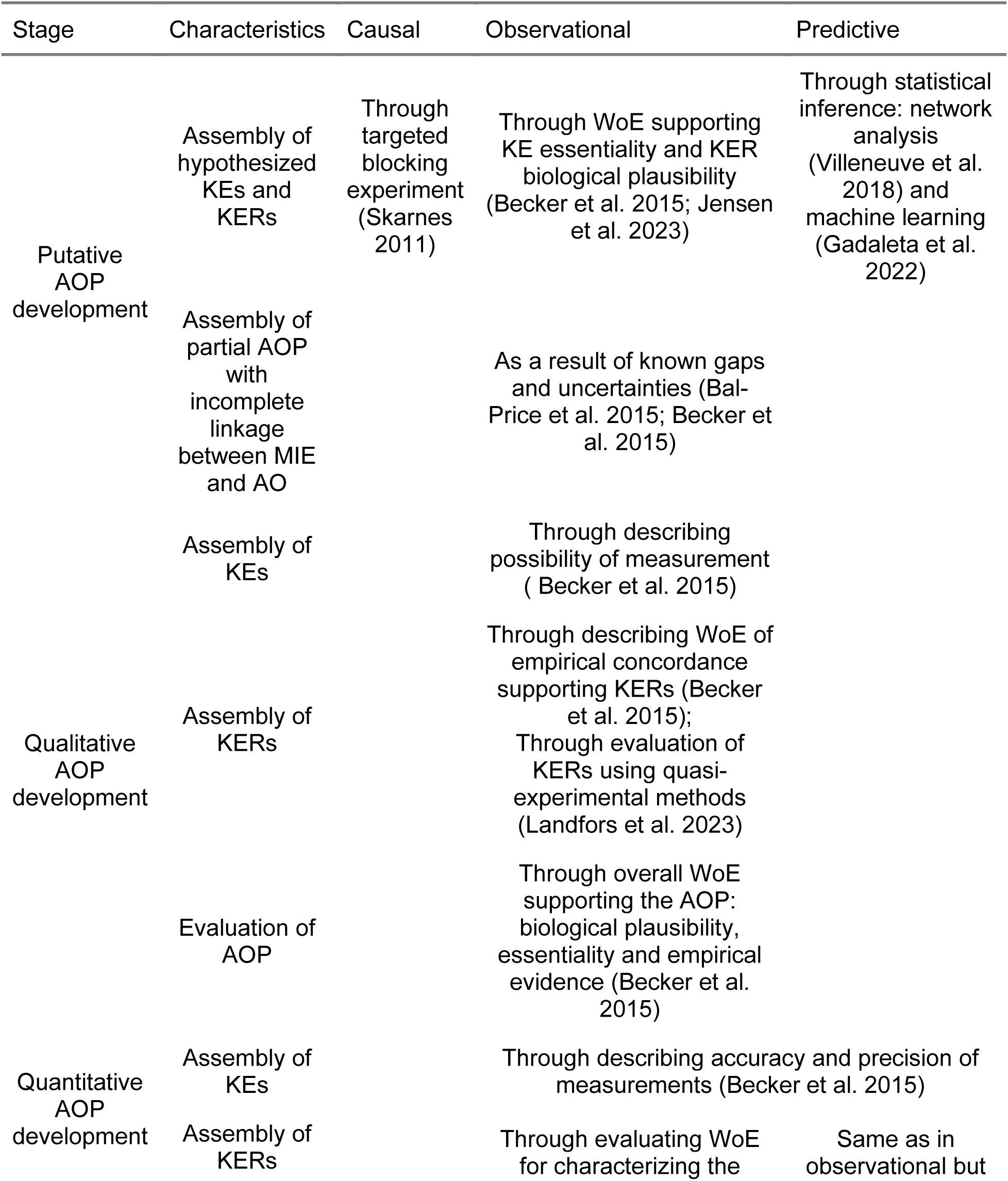
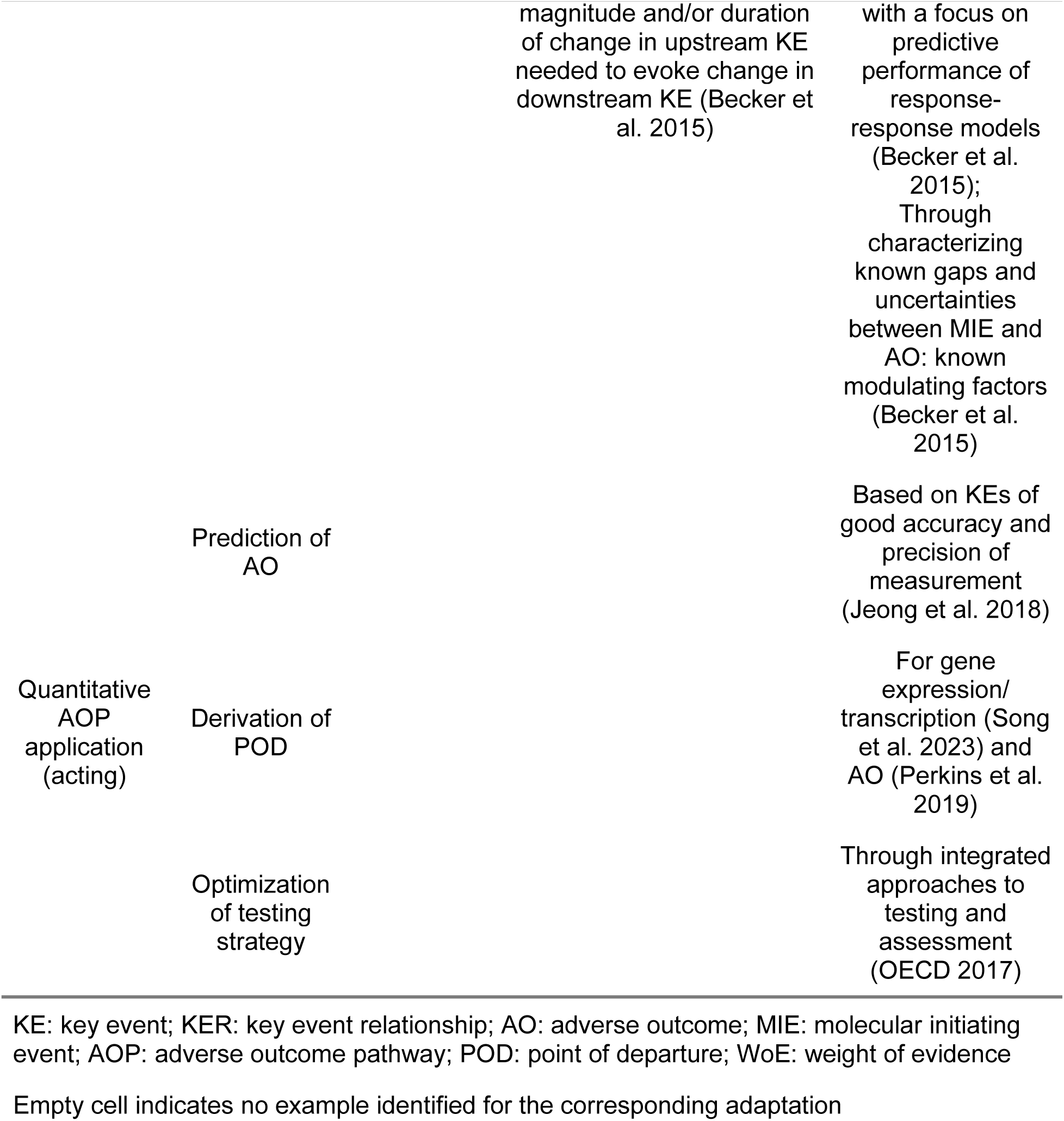
Examples of adaptation of KER terminology in AOPs and qAOPs from published literature.

Adapting the term “causal” in the strict sense would require experiments implementing direct randomization of upstream KEs. However, no examples of practical implementation could be identified in the literature. The alternative approach in published studies is targeted blocking experiments to infer the impact of *KE*_*up*_ on *KE*_*down*_ that a reduction or complete absence of a *KE*_*down*_ when *KE*_*up*_ is blocked (Meek et al. 2014). Empirical evidence from targeted blocking experiments are direct support and contribute to a high level of confidence for evaluating the essentiality of KEs (Meek et al. 2014; Villeneuve et al. 2014b; Villeneuve et al. 2024). The development and application of qAOPs often assumes that KEs are essential (Jensen et al. 2023) and KERs are causal (Burgoon et al. 2020; Gou et al. 2023).

Observational analyses of KEs, KERs and overall AOP could involve weight-of-evidence (WoE) assessments of supporting evidence and quasi-experimental assessments. The evolved MOA Hill considerations (Meek et al. 2014) are adapted in the handbook and the OECD guidance for the WoE assessment of the essentiality of KEs, the biological plausibility of KERs, the concordance of KER empirical evidence and the overall AOP level of confidence (OECD 2017; Villeneuve et al. 2024). Best practice has been recommended for the adapted WoE assessments (Villeneuve et al. 2014b) to categorize confidence levels as high, moderate or low (Becker et al. 2015). Characterizing known gaps and uncertainties that modulate incomplete linkages between MIE and AO could further enhance the weights of evidence in the overall AOP (Bal-Price et al. 2015; Villeneuve et al. 2024). When high WoE is achieved, qualitative and quantitative descriptions of KE measurement methods and their accuracy and precision are crucial (Villeneuve et al. 2014b). These descriptions support the characterization and calibration of predictive modeling of relationships among the sequence of KEs in the overall AOP (Becker et al. 2015). In addition to WoE assessments, the instrumental variable method, a type of quasi-experimental design, shows promise in providing strong evidence supporting KER causality (Landfors et al. 2023).

Predictive models are primarily involved in the putative and quantitative stages (OECD 2017). Assembly of KEs and KERs in the putative stage can be derived using machine learning (Gou et al. 2023; Hemmerich and Ecker 2020) or probabilistic graphical network models (Villeneuve et al. 2018). KEs could be screened from a pool of candidate molecular and cellular events using feature selection techniques (Perkins et al. 2019). Similar methods have been applied to screen for genes and genomic modules associated with significant differential gene expressions (Callegaro et al. 2021; Hemmerich and Ecker 2020). Topological metrics are evaluated to identify nodes and edges that best improve overall predictive performance on the AO as KEs and KERs, respectively (Villeneuve et al. 2018). Graphical network models might consist of deterministic and probabilistic nodes, and undirected, directed, acyclic edges, that have been involved in existing studies of AOP development and application (OECD 2017; Perkins et al. 2019; Villeneuve et al. 2018).

Quantitative AOP development focuses on the derivation of response-response models [Villeneuve2024]. Studies of qAOP typically assume the causality of KERs has been established (Burgoon et al. 2020; Gou et al. 2023; Jensen et al. 2023), supported by the essentiality of KEs, the biological plausibility of KERs and the concordance in KER empirical evidence. Furthermore, to assure the reliability of qAOPs within relevant applicability domains, the handbook includes a guidance (annex 2) to evaluate the WoE of qAOPs by not only predictive performance, also considerations for known modulating factors, feedback loops and generalizability across the applicability domains. Examples of possible regulatory application of qAOPs include prediction of AO given MIE (Jeong et al. 2018), derivation of omics (Song et al. 2023) and *in vitro* (Perkins et al. 2019) PODs, and optimization of testing strategies (OECD 2017).

In summary, the handbook’s nuanced usage of terms is reflected in AOP development that focus of adaptation varies across stages of AOP development. Given the challenge of obtaining experimental evidence for achieving causality, targeted block experiments offer a practical approach to balance weights of evidence and feasibility. Best practice (Becker et al. 2015; Villeneuve et al. 2014b) has been recommended to implement observational inference using WoE assessments based on the evolved MOA Hill considerations (Meek et al. 2014). The stage of putative AOP development has a dual focus on observational and predictive evaluations. The qualitative stage focuses on the WoE of evidence supporting KEs, KERs and the overall AOP. The stage of qAOP application and application involves various predictive approaches, including machine learning, network and response-response models.

## 4 DISCUSSIONS

The nuanced terminology usage in the handbook and approaches of adaptation reflect the complex nature of AOP development. Our results suggest several issues that may warrant further discussion, including: 1) a mismatch between the terminology in guideline and the methodology in practice, and 2) considerations of terminology for interdisciplinary communication.

### 4.1 Mismatch between methodology and terminology

Some putative AOP studies inferred causality of KERs based on predictive models (Gou et al. 2023; Jeong et al. 2018). We argue the use of predictive models in these studies supported the predictive but not the causal aspect of KERs, highlighting a mismatch between the treatment effect framework and predictive models. This mismatch could be demonstrated through possible misinterpretations of probabilistic network models as causal. Probabilistic network models (Koller and Friedman 2009) are based on directed acyclic graphs (DAGs). DAGs are commonly involved in causal reasoning and designing of structural equation modeling (Greenland et al. 1999), e.g. to approximate unbiasedness by quasi-experimental methods. But the use of a DAG is not a guarantee for having a causal model. It depends on context, if a model involving DAGs can provide unbiased average treatment effects for causal inference. In Bayesian modelling, Directed acyclic graphs are used to decompose the joint probability distribution of variables and parameters into a product, with no implication of causality. A Bayesian network is usually referred to a probabilistic network model with binary or categorical nodes. Bayesian networks comprise a wide range of applications, where the network can be inferred from multivariate data, e.g. in machine learning, or given as representative of causal relationships where the probabilities or conditional probability distribution are informed by expert judgment, e.g. in decision analysis. Bayesian networks have been applied to parameterize models where confounding biases are adjusted using quasi-experimental methods such as the backdoor criterion (Pearl 2009) and instrumental variables (Shingaki et al. 2021). With the popularity of Bayesian networks (Kaikkonen et al. 2021; Spinu et al. 2020) comes misconceptions of to what extent a Bayesian network indicates causality, which might come from the use of a DAG. There are studies where results of a Bayesian network were interpreted as causal without employing bias reduction (Chen and Pollino 2012; Jeong et al. 2018; Zou et al. 2010). For evaluating the uncertainty in a Bayesian network, beyond the inherent uncertainty embedded within the probability model, it is crucial to consider the process and context in which the Bayesian network is applied (Sahlin et al. 2021).

This mismatch is particularly crucial in the stage of putative AOP development, where the assembly of KEs and KERs could be achieved through either causal inference or statistical modeling. The reason is, although predictive inference of KEs and KERs is optional in the recommended best practice for the putative stage (Villeneuve et al. 2014b), an AOP would be incomplete without considerations on the essentiality of KEs, biological plausibility and empirical support of KERs. Therefore, misrepresenting results from predictive models as causal may falsely imply that the dual requirements have been fulfilled in the putative stage, which could risk overlooking causal and observational assessment in the qualitative stage.

Furthermore, this mismatch also highlights the importance of integrating predictive models with the evidence from causal or observational inference in the qAOP stage. Current qAOP development often assumes that causality of KERs is well-supported in established AOPs (Burgoon et al. 2020; Gou et al. 2023; Jensen et al. 2023). Yet there are limited approaches to propagate the assumed causality in predictive models. One potential approach is the use of Bayesian methods to integrate known causality of KERs and prior knowledge on KEs in qAOP (Spînu et al. 2022). Another approach involves incorporating causal KERs on the target/ utility/ loss function of predictive models through regularization techniques such as Lasso (Tibshirani 1996). While regularization has been applied in in silico model to mitigate overfitting (Hemmerich and Ecker 2020), there has been limited exploration of its use in adjusting the target/ utility/ loss function on known causality. This may be due to concerns that regularizing on causality may penalize predictive performance. However, it is crucial to re-emphasize that AOPs are only complete with both causal and predictive characteristics (Becker et al. 2015) and good predictive performance alone is insufficient for robust qAOP development. According to the handbook’s guideline on the WoE assessment of qAOPs (annex 2), known modulating factors and feedback loops need to be considered along with predictive performance for a qAOP to achieve high level of confidence (Villeneuve et al. 2024).

### 4.2 Careful considerations of terminology in AOP guidance

We would like to initiate a discussion for careful considerations on the usage of terms “causality” and “essentiality” to facilitate interdisciplinary communication and collaboration.

The use of term “causal” needs to be considered because the frameworks for evaluating causality in the handbook (Villeneuve et al. 2024) and for the use in integrated testing strategies (OECD 2017) may not necessarily conclude causality in the same sense employed in toxicology and generic scientific reasoning. Even the recommended best practice (Becker et al. 2015; Villeneuve et al. 2014b) may be insufficient to achieve unbiased estimates of KE essentiality and KER causality. This is not a critique of current established AOP guidelines and practice because, as shown by our results, the term “causal” is pronounced and utilized in a consistent manner by the handbook, the OECD guidance and studies within the niche of AOP development. Distinctive usage across fields of the same terminology is common.

However, adding clarifying annotations on the term “causal” could greatly facilitate interdisciplinary communication and collaboration that potential bias may exist and that the results could be “causal estimates” rather than “causal”.

To demonstrate, the handbook’s framework highlights the inclusion of direct and indirect evidence and a prioritization of direct evidence in the assessment of essentiality of KEs and biological plausibility of KERs. Direct evidence of KE essentiality (handbook section 4B, Table 5) includes evidence from targeted blocking experiments, such as know-out experiments (Skarnes et al. 2011). Knockout experiments pragmatically approximates counterfactual inference and are more robust and reliable than observational evaluation of multiple KEs (see supplementary information). However, results from targeted blocking experiments are not always causal because the caveat of off-target issues applies in design (Höijer et al. 2022). The instrumentation of blocking could be attributable to confounding and uncontrolled mediation on downstream KEs (Skarnes et al. 2011). Unbiased causality could be achieved only if the blocking methods induce blocking in the target *KE*_*up*_ but not anything else that causes changes in *KE*_*down*_ or their modulating factors. If the instrument to block *KE*_*up*_ activates a third, unobserved event that, e.g., increases the activity of *KE*_*down*_, it is possible to observe no change in *KE*_*down*_, because the reduction by blocking *KE*_*up*_ could be canceled out by the activation from the third event.

In targeted knockout experiments, there could be at least two technical reasons leading to the off-target issue. Firstly, the genetic target in knockout procedure is identified by genome-wide association studies, which is association by definition. Confounders may exist since the gene-protein associations are not always one-to-one matched (Olariu et al. 2021). It is well known that the same protein can be coded by different genes or multiple copies of the same gene at different loci (Campenhout et al. 2019). It is also not uncommon to observe the synthesis of multiple proteins from the transcription of one gene. The variational splicing of introns of one gene can be transcripted to different mRNAs, which result in the production of multiple, distinctive proteins (De Conti et al. 2013). Targeted mutation techniques have the potential to become quasi-experimental by implementing bias adjustment methods such as instrumental variables (Landfors et al. 2023). Secondly, assuming the target identification is free from confounding bias, unintended genetic changes could be induced during gene editing (Höijer et al. 2022). Therefore, direct evidence from targeted blocking experiments, as prescribed in the best practice, could be insufficient to guarantee causality.

Indirect, non-experimental evidence is evaluated using a WoE framework based on the evolved MOA Hill considerations (Meek et al. 2014). The evolved MOA Hill considerations classifies weights of confidence into high, moderate or low rather than a dichotomous judgement of causality. This is evident that the terms “causal”, “causality” or etc. were included only once (not including quotes) in the source paper (Meek et al. 2014) to describe the consideration for KE essentiality that *“if following cessation of repeated exposure for various periods, effects are reverse, this constitutes relatively strong evidence that key events are causal”*. Furthermore, the adaptations made by the handbook implement no further bias adjustment and focus on the classification of WoE of supporting evidence.

In conclusion, the frameworks for KER evaluation in the handbook and recommended practice could be insufficient to reach the causal requirement prescribed in the handbook.

We also would like to bring up a discussion on the use of the term “essential”, particularly in the essentiality of KEs. We argue that adding enhancing annotations on the term “essential” could benefit its practical utility in the assessment of KE essentiality. It is crucial to note that this discussion does not revolve around the distinction between mathematical and toxicological context of “essential”. The term “essential” has specialized mathematical context and usage to describe the conditional relationship between two statements. To claim mathematical essentiality of event A for event B, that is to say, if A does not occur, then B cannot occur at all. The validation of mathematical essentiality requires a falsity test. Any case where the occurrence of B without the occurrence of A falsifies essentiality.

Obviously, such a falsity test is practically difficult to perform in science (Graesser and Hu 2011), especially in biology and toxicology. Most biological events are the consequence of one or several alternative, sufficient causes. The use of “essential” in toxicology is distinct from the mathematical context, including the evolved MOA Hill considerations and AOP development. Under the toxicological context, evaluation of essentiality considers the impact of blocking or reversing *KE*_*up*_ on *KE*_*down*_ (Meek et al. 2014). A mathematically essential event would cause complete absence in the consequence, while a toxicologically essential event could describe a *KE*_*up*_ which, when blocked, prevent or significantly modify *KE*_*down*_ (Meek et al. 2014; Villeneuve et al. 2024). The validation of mathematical essentiality of *KE*_*up*_ for *KE*_*down*_ could be practically unfeasible. The guidelines for AOP development needs to be a pragmatic framework that balances scientific rigor and practicability to serve regulatory and industrial application in the real world. However, the level of significance of modification/ reduction is subjective. Neither the evolved MOA Hill considerations nor the handbook defines the level of modification in *KE*_*down*_ to which the changes associated with blocking *KE*_*up*_ should be considered as significant and sufficient. Furthermore, the level of modification/ reduction is diverse across systems, organs and molecules by nature. For example, body weight change depends on a myriad of factors, including time period, age, sex and baseline weight. It would be helpful to annotate the handbook’s defining question of KE essentiality by adding a description of the level at which the modification in *KE*_*down*_ is both statistically significant and biologically sufficient. Further research using the benchmark dose modeling methodology might be useful to characterize the level of modification for KE essentiality.

## 5 CONCLUSIONS

This study aimed to explore the usage and adaptation of the terms “causal”, “observational” and “predictive” for KER description in AOP guidance documents. The results indicate that KERs are described to fulfill both causal and predictive requirements across three stages of AOP development, and distinct approaches are implemented to address different focuses of adaptation across these stages. These findings suggest that the nuanced approach towards the distinction between causal and predictive inference of KERs reveal a mismatch between predictive models and causality in published studies of AOP development, particularly pertaining to the putative stage. Further exploration is needed to integrate causality in qAOP development and application. Further discussions could be helpful to consider the handbook’s usage of the terms “causal” and “essential” to facilitate more comprehensive, transparent interdisciplinary communication and practical application of AOP under the academic, regulatory and industrial contexts.

