## Supplementary Information for "Causal, Predictive or Observational? Different Understandings of Key Event Relationships for Adverse Outcome Pathways and Their Implications on Practice"

### 1 List of Definitions and Abbreviations of Key Terms in This Paper

*Table S1.1: Abbreviations of Important Terms in This Paper*

| Abbreviation | Full Terminology |
|---|---|
| 3R | Reduce, Refine and Replace |
| OECD | Organisation for Economic Co-operation and Development |
| ASPIS | Animal-free Safety assessment of chemicals: Project cluster for Implementation of novel Strategies |
| NAM | Non-animal Methodology |
| QST | Quantitative System Toxicology |
| MOA | Mode of Action |
| AOP | Adverse Outcome Pathway |
| qAOP | Quantitative Adverse Outcome Pathway |
| MIE | Molecular Initiating Event |
| KE | Key Event |
| KER | Key Event Relationship |
| AO | Adverse Outcome |
| RCT | Randomized Controlled Trial |
| WOE | Weight of Evidence |
| MLE | Maximum Likelihood Estimation |
| DAG | Directed Acyclic Graph |
| BN | Bayesian Network |

*Table S1.2: Glossary of Key Terms in This Paper*

| Terminology | Definition | Source |
|---|---|---|
| Adverse Outcome Pathway | An AOP describes a | Villeneuve et al. (2024) |

| Terminology | Definition | Source |
| --- | --- | --- |
| | sequence of events commencing with initial interaction(s) of a stressor with a biomolecule within an organism that causes a perturbation in its biology (i.e., molecular initiating event, MIE), which can progress through a dependent series of intermediate key events (KEs) and culminate in an adverse outcome (AO) considered relevant to risk assessment or regulatory decision-making. | |
| Molecular Initiating Event | A specialised type of key event that represents the initial point of chemical/stressor interaction at the molecular level within the organism that results in a perturbation that starts the AOP. | Villeneuve et al. (2024) Table 1 |
| Key Event | A change in biological or physiological state that is both measurable and essential to the progression of a defined biological | Villeneuve et al. (2024) Table 1 |

| Terminology | Definition | Source |
| --- | --- | --- |
| | perturbation leading to a specific adverse outcome. | |
| Key Event Relationship | A scientifically-based relationship that connects one key event to another, defines a causal and predictive relationship between the upstream and downstream event, and thereby facilitates inference or extrapolation of the state of the downstream key event from the known, measured, or predicted state of the upstream key event. | Villeneuve et al. (2024) Table 1 |
| Adverse Outcome | A specialised type of key event that is generally accepted as being of regulatory significance on the basis of correspondence to an established protection goal or equivalence to an apical endpoint in an accepted regulatory guideline toxicity test. | Villeneuve et al. (2024) Table 1 |

| Terminology | Definition | Source |
|---|---|---|
| Randomized Controlled Trials | A randomized controlled trial is a prospective, comparative, quantitative study/experiment performed under controlled conditions with random allocation of interventions to comparison groups | Bhide et al. (2018) |
| Causal Inference | Causal inference pursues the unbiased understanding of how one condition, the cause, affects another condition, the outcome, free from the threats to internal validity. | Guzelian et al. (2005) |
| Observational Inference | Observational inference refers to the inference of causality based on data that is obtained through the process of observation without randomized experimentation. | Campbell and Stanley (2015) |
| Quasi-experiment | A quasi-experiment is an empirical interventional study used to estimate the causal impact of | Campbell and Stanley (2015) |

| Terminology | Definition | Source |
|---|---|---|
| | an intervention on target population without random assignment | |
| Predictive modeling | (Predictive models have) the goal of maximizing the predictive accuracy of the outcome equation. | Young (2019) |
| Mode of Action | A postulated mode of action (MOA) is a biologically plausible sequence of key events leading to an observed effect supported by robust experimental observations and mechanistic data. | Meek et al. (2003) |
| Point of Departure | The dose-response point that marks the beginning of a low-dose extrapolation. This point can be the lower bound on dose for an estimated incidence or a change in response level from a dose-response model (BMD), or a NOAEL or LOAEL for an observed incidence, or | U.S. EPA (2022) |

| Terminology | Definition | Source |
|---|---|---|
| | change in level of response. | |

The definitions of KE and KER presented in this table correspond to the first appearance of them in INTRODUCTION of the handbook. There are various, distinctive definitions of these terms as outlined in the main text.

### 2 EXAMPLES ON THE DIFFERENCE BETWEEN PREDICTIVE AND CAUSAL RELATIONSHIPS

The first example is to demonstrate statistical correlation is not always causation. It has been commonly used in pedagogy regarding the association between ice cream sales and drowning incidents over seasons. There is a strong correlation between ice cream sales and drowning incidents because both increase in summer. However, there is no causal relationship between ice cream sales and drowning. Both are consequences of the increase in temperature in summer, which is a confounding variable.

The following example demonstrates that causality can exist between two events of statistical independence. Imagine driving a normally functional motor vehicle on a hilly road. The accelerator pedal controls the speed of the vehicle. During the uphill segment, the driver presses the pedal to maintain the speed at 60 km/h. During the downhill segment, the driver releases the pedal to also maintain the speed at 60 km/h. Since the speed is held at a constant up and down hill, there is zero correlation between speed and the pedal movement.

### 3 RANDOM ASSIGNMENT OF TREATMENT STATUS APPROXIMATES UNBIASED ESTIMATE OF TREATMENT EFFECTS

To understand the difference between observational and experimental analysis, it is important to review the gold standard of toxicological study, i.e., randomized controlled trials (RCT) (James et al. 2015).

Causal reasoning aims for an unbiased estimation of the treatment effect on the outcome. The unbiasedness is achieved through the isolation of the treatment effect from all other observed and unobserved factors (Cinelli et al. 2022). Such isolation requires the evaluation of counterfactual, which is not feasible given current human technology (Lipton and Ødegaard 2005). Randomized controlled trials is the best approximation to counterfactual for unbiased estimates of average treatment effects (Lipton and Ødegaard 2005). This is achieved by trading external validity for internal validity of the results (Rothwell 2005).

Subjects are randomized to receive different treatment assignments in a controlled laboratory environment. Consider the simple example of one control group and one treatment group. The randomization process ensures that the only difference between the two groups is the treatment status. All other observed and unobserved factors that may affect either the treatment process and/or the outcome are balanced on average between the two groups (Cinelli et al. 2022). All the potential threats to internal validity are eliminated through randomization. Therefore, the average difference in the outcome between the two groups after treatment is an unbiased estimate of the treatment's causal impact on the outcome (Angrist and Pischke 2009).

A typical toxicological experiment expands this simple example design to compare multiple intervention arms or dose levels. In other words, subjects are randomly assigned to different experimental groups. The average difference between the control and each arm/dose group is an unbiased estimate on the outcome specific to that experimental group.

In summary, the essence of experimentation is the random assignment of treatment status.

### 4 TARGETED MUTATION ENHANCES ACCURACY AND REDUCES BIAS IN KER ANALYSIS

Technologies of targeted mutation, such as gene knockout (Skarnes et al. 2011) and genome editing (CRISPR-Cas9 (Campenhout et al. 2019)), can greatly mitigate the biases in the estimated impact of targeted genes on observed phenotypes. The targeted genes can be linked with proteins using genome-wide association studies (Tam et al. 2019). KEs can be identified when the proteins associated with targeted mutations are major contributors to the observed phenotypes. For instance, the arsenic methyltransferse (AS3MT) protein has been hypothesized to play a major role in the toxicity of inorganic arsenic through methylation (Cohen et al. 2006; Stýblo et al. 2002). Knockout mice are created from embryonic cells with targeted mutations inserted to the DNA fragments which are statistically associated with AS3MT protein (Drobna et al. 2009). Compared with their wild types, knockout mice shows much more significant metabolic phenotypes, such as fat accumulation, insulin resistance and body mass increase in RCTs (Douillet et al. 2017). This association support the protein AS3MT as a KE for metabolic phenotypes in mice.
